# NetGO: Improving Large-scale Protein Function Prediction with Massive Network Information

**DOI:** 10.1101/439554

**Authors:** Ronghui You, Shuwei Yao, Xiaodi Huang, Fengzhu Sun, Hiroshi Mamitsuka, Shanfeng Zhu

## Abstract

Automated function prediction (AFP) of proteins is of great significance in biology. In essence, AFP is a large-scale multi-label classification over pairs of proteins and GO terms. Existing AFP approaches, however, have their limitations on both sides of proteins and GO terms. Using various sequence information and the robust learning to rank (LTR) framework, we have developed GOLabeler, a state-of-the-art approach of CAFA3, which overcomes the limitation of the GO term side, such as imbalanced GO terms. Unfortunately, for the protein side issue, available abundant protein information, except for sequences, have not been effectively used for large-scale AFP in CAFA. We propose NetGO that is able to improve large-scale AFP with massive network information. The novelties of NetGO have threefold in using network information: 1) the powerful LTR framework of NetGO efficiently and effectively integrates both sequence and network information, which can easily make large-scale AFP; 2) NetGO can use whole and massive network information of all species (>2000) in STRING (other than only high confidence links and/or some specific species); and 3) NetGO can still use network information to annotate a protein by homology transfer even if it is not covered in STRING. Under numerous experimental settings, we examined the performance of NetGO, such as general performance comparison, species-specific prediction, and prediction on difficult proteins, by using training and test data separated by time-delayed settings of CAFA. Experimental results have clearly demonstrated that NetGO outperforms GOLabeler, DeepGO, and other compared baseline methods significantly. In addition, several interesting findings from our experiments on NetGO would be useful for future AFP research.

## 1 Introduction

As the most basic structural molecule, protein maintains the basic cell activities and biodiversity [1]. Identifying protein/gene functions is of great significance to understand the nature of biology. For this purpose, gene ontology (GO) was launched in 1998, currently becoming the most influential ontology [2]. So far, GO contains 49,177 biological concepts (Nov. 2017), covering three different domains of Molecular Functional Ontology (MFO), Biological Process Ontology (BPO) and Cellular Component Ontology (CCO). Due to the advancement of sequencing technologies, the number of available protein sequences has been explosively increased. Only a very tiny part of newly obtained sequences, however, have experimental GO annotations. For example, only less than 0.2% of around 112 million protein sequences in UniProKB (April 2018) have experimental GO annotations [3]. This is because identifying protein experiments is both time- and resource-consuming. As such, automated function prediction (AFP) has become increasingly important in reducing the gap between the huge number of protein sequences and very limited experimental annotations functions by biological[4, 5].

For advancing the research of large-scale AFP, a competition called Critical Assessment of Functional Annotation (CAFA)^1^ has been held three times, i.e. CAFA1 in 2010-2011, CAFA2 in 2013-2014 and CAFA3 in 2016-2017 [4, 5]. CAFA used a time-delayed evaluation procedure to assess the accuracy of protein function prediction submitted by participants. A large set of target proteins (around 100,000 in CAFA2 and CAFA3) were first available to the participants, who were required to submit their predictions before the deadline (T0). A few months later (T1), target proteins with experimental annotations were then used as benchmark for performance evaluation. In CAFA2, the benchmark data was divided further into two categories: *no-knowledge* and *limited-knowledge*. The *no-knowledge* benchmark proteins are those with no experimental annotations before T0, and at least one experimental annotations before T1. The *limited-knowledge* benchmark proteins are those with the first experimental annotations in the target domain between T0 and T1 and experimental annotations in at least one other domain before T0. Currently more than 99% of all proteins have no experimental annotations. Therefore our focus in this study is on *no-knowledge* benchmark.

From a machine learning viewpoint, AFP is a large-scale multilabel classification problem, where multiple GO terms (labels) can be assigned to each protein (instance) [6]. AFP faces two main challenges on the GO (label) and protein (instance) sides. On the GO side, each protein can be assigned to multiple GO terms, because GO is a hierarchical structure. If a protein is assigned by a GO term, for example, all GO terms at ancestor nodes (in GO) of this term are assigned to this protein as well. The experimental GO annotations of human proteins in Swissprot [7] (Dec. 2017) reveal that each human protein is annotated by 74 GO terms on average. On the protein side, information about proteins are not limited to sequences. Sequences are just part of all information about proteins. In particular, sequences are static and genetic information, while proteins are alive, and dynamic. In addition, environmental information might affect protein activity more than static, genetic and inherited information. Thus an imperative issue is how to use multiple types of data other than protein sequences for AFP.

The results of past CAFA on the *no-knowledge* benchmark show that sequence-based AFP (SAFP) methods can be the best prediction methods. Even simple homology-based methods, which use BLAST or PSI-BLAST, are very competitive [8, 9, 10]. For instance, [10] shows that their methods implementing simple homology-based inference performed only slightly worse than the best method by the Jones-UCL group in CAFA1. Recently we developed a high-performance SAFP method, GOLabeler [11]. The initial evaluation on CAFA3^2^ revealed that GOLabeler achieved the first place out of nearly 200 submissions by around 50 groups all over the world in terms of F_max_ (see Section 4.2) in all three GO domains. GOLabeler addressed the challenge of the label side, by using multiple types of sequence-based evidences such as homology, domain, family, motif, amino acid k-mer, as well as biophysical properties. GOLabeler seamlessly integrated these evidences in the “learning to rank” (LTR) framework[12], which is especially effective and efficient for tackling the large-scale multilabel learning problem [13, 14, 15]. LTR is widely used in ranking web pages with respect to a query, while it is used in GOLabeler for ranking GO terms with respect to a protein (See Supplement and [11] for more introduction on LTR). As a state-of-the-art, large-scale AFP method in CAFA settings, GOLabeler uses sequence information only. However, many protein functions cannot be easily inferred from protein sequences only. For example, a well-accepted hypothesis of network-based methods is that interacting proteins should share similar functions under the “guilt by association” (GBA) principle [16, 17]. A reasonable question is to explore other types of protein information to improve the performance of GOLabeler or more generally AFP further.

We propose a new AFP method called NetGO. The basic idea of NetGO is to incorporate the network-based evidence into the GOLabeler framework (i.e. LTR) by which to improve the performance of a large-scale AFP. In other words, NetGO addresses the both two sides of the challenges: 1) the label side of multilabel classification problem by using LTR and 2) the instance (protein) side by incorporating network-based information. More importantly, our method makes it feasible to incorporate network information at a large-scale level. The performance of NetGO has been thoroughly validated by conducting comprehensive experiments on large-scale datasets under the CAFA settings. We compared NetGO with GOLabeler, a state-of-the-art method. Experimental results indicate that NetGO has significantly outperformed GOLabeler in both BPO and CCO. This validates the effectiveness of network-based evidence to improve the performance of predicting GO terms in BPO and CCO. In addition, we came across a variety of interesting discoveries in our experiments. For example, in the networks we used, the proteins that can improve the BPO prediction tend to have higher node degrees (more weighted connections) than those which cannot. However this difference was not found in MFO. Another interesting finding was that the performance had been improved by NetGO over GOLabeler against all groups of GO terms (appeared more than 10 times) in both BPO and CCO. This means that the improvement was not contributed by only part of GO terms. Finally, we report a typical example by applying NetGO. This example has demonstrated that NetGO outperformed GOLabeler as well as other baseline methods in that it had correctly predicted the largest number of all GO terms with the fewest errors.

## 2 Related work

For AFP, not only sequences but also various biological information, such as genomic context, gene expression and biomedical literature can be used [18]. In this work, we focus on network information for AFP, where each node in a network represents a protein and each edge indicates the interaction between two corresponding nodes (proteins). The standard implementation of GBA for Network-based AFP is to predict the function of a target by using the functional labels of its direct neighbors in the same network, such as majority voting [19].

Many studies have been conducted by using network data for AFP. Most of them deal with only proteins of specific species and/or a limited number of GO terms. For example, [20] used Markov random field (MRF) for function prediction of proteins in Yeast over 134 GO terms in BPO. GeneMANIA [21] and ClusDCA [22] are two state-of-the-art network-based general AFP methods. GeneMANIA used label propagation over an integrated network, starting with mouse, and then extending to multiple organisms, such as human and yeast. ClusDCA is unique in the sense that embedding of protein and GO terms is trained from different graphs, i.e. protein networks and GO ontology, respectively. A notable weakness of these two approaches is that they train the prediction model of each species independently. As a result, they cannot annotate proteins that are not covered in the network. Moreover, sequence information are completely ignored. More recently ProSNet was proposed to integrate both sequence homology and molecular network information of 5 species for constructing a large heterogeneous network to improve the performance of AFP [23]. Due to its high complexity in constructing and training a global heterogeneous network, it would be infeasible for ProSNet to incorporate the network of hundreds of species or more at the same time. In addition, the performance of ProSNet was examined by cross-validation, which, in the network, cannot clearly separate test data from training data. This casts doubt on whether predicting real “new” proteins is implemented in the cross-validation. Note that, for real “new” proteins, a lot of (true) edges would be unknown in the network. In contrast, our performance validation uses the CAFA setting, where the benchmark test data is a large number of *no-knowledge* proteins in various organisms. This experimental procedure allows our experiments to avoid the above issue of cross-validation.

There already exist several attempts on using network information for AFP under the CAFA setting with limited success. For example, MS-kNN, a top method in both CAFA1 and CAFA2, used a weighted average method to integrate multiple sources (information of each source through *k*-nearest neighbor (KNN)) including networks [24]. However, the accuracy by MS-kNN was similar to that by KNN from a single source, i.e. sequence similarity, probably because of the limited input data. DeepGO used deep learning to learn features of proteins from both sequences and networks in CAFA3 [25]. DeepGO focused on a limited number of GO terms, due to computational limitation and sparse training data. DeepGO improved the performance by adding network information to sequence information only under cross-validation, by which still the effect of incorporating network information under the CAFA setting is unclear. GAS-C (Guilt by Association with Consensus) used a consensus approach to integrate network information with BLAST[26]. Training data were 10,000 experimentally annotated targets in SwissProt and the network was with binary edges with highest confidence from STRING [27]. GAS-C was better than both BLAST and GAS under all domains, but not than even Naive in MFO. COFACTOR also used a consensus approach to integrating three types of information [28]. However it is not shown that incorporating the network information is effective. Overall, the preliminary results of CAFA3 for the above methods show that incorporating network information effectively for better AFP is still a challenging problem.

Differing from the above models, NetGO uses network information in three novel ways: 1) The powerful LTR framework of NetGO efficiently and effectively integrates both sequence and network information, which can easily make large-scale protein function prediction. 2) NetGO uses whole and massive network information in STRING (other than only high confidence links and/or some specific species). 3) NetGO can effectively annotate proteins by homology transfer in the absence of network information.

## 3 Methods

### 3.1 Notation

Let *D* be given training data with *N*_*D*_ proteins, i.e. |*D*| = *N*_*D*_. Let *G*_*i*_ be the *i*-th GO term, and 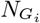 be the number of proteins with *G*_*i*_ in *D* (Note that this number is obtained by considering the structure of GO. That is, if *G*_*i*_ is assigned to a protein, this protein is with all GO terms of the ancestors of *G*_*i*_ in GO.). Let *T* be the given test data (the number of proteins: *N*_*T*_ = |*T*|), in which let *P*_*j*_ be the *j*-th protein. Let *I*(*G*_*i*_, *p*) be a binary indicator, showing if *G*_*i*_ is the ground-truth (true) GO term of *p*. That is, *I*(*G*_*i*_, *p*) is one if true; otherwise zero. Let *S*(*G*_*i*_, *P*_*j*_) be the score (obtained by a method), showing that *P*_*j*_ is with *G*_*i*_, where in ensemble methods, *S*_*k*_(*G*_*i*_, *P*_*j*_) is the predicted score by the *k*-th component method.

For a given organism, we have *m* types of protein networks *PN*^(*l*)^ (*l* = 1*,…,m*) from different sources, such as genomic context, physical interaction and database. Each network *PN*^(*l*)^ consists of a set of nodes *PV*^(*l*)^ and a set of edges *PE*^(*l*)^ between a pair of two nodes. Each node corresponds to a protein in this organism, and an edge represents a possible interaction (association) between two proteins. Let *PE*^(*l*)^(*i*, *j*) be the edge between two nodes 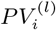 and 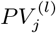, and *ω*^(*l*)^(*i*,*j*) be the weight of edge *PE*^(*l*)^(*i*, *j*) ∈ [0, 1] measuring the confidence of association between two nodes 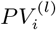 and 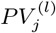. Given target protein *P*_*j*_ and *i j* GO term *G*_*i*_, the core idea of NetGO is to estimate *S*(*G*_*i*_, *P*_*j*_) by using all available networks *PN*^(*l*)^(*l* = 1…,*m*) of different organisms.

### 3.2 NetGO: Overview

Fig 1 shows the entire framework of NetGO for AFP. In testing, given the sequence of a query protein, candidate GO terms are generated from six components, which are already trained by using different information. Among the six components, five components were originally developed in GOLabeler (so they are from sequence information), while the rest one component, which we call Net-KNN, is from network information and newly developed in this study. Each candidate GO term receives prediction scores from the six components, resulting in a feature vector of length six. Then candidate GO terms, i.e. feature vectors, are the input into the learning to rank (LTR) model, which is also already trained by using training data. Finally, a ranked list of GO terms is returned as the final output of NetGO.

### 3.3 NetGO: Five components from GOLabeler

GOLabeler [11] is an ensemble model by LTR, with five component methods from protein sequence information: Naive, BLAST-KNN, LR-3mer, LR-Interpro and LR-ProFET (Note: KNN and LR stand for *k*-nearest neighbors and logistic regression, respectively). These five components are different and complement to each other. Before giving more detailed description of Naive and BLAST-KNN, we briefly introduce the other three LR-based components using different features: 1) LR-3mer: we count the frequency of amino acid trigram (3mer) in each protein resulting in 8,000(= 20^3^) features. 2) LR-InterPro: we run InterProScan^3^ to get 33,879 binary features that represent the occurrence of a large number of motifs, protein families and domains in InterPro [29]. 3) LR-ProFET: ProFET [30] has been used in various function prediction task, which consists of 1170 features including elementary biophysical properties and local potential features etc.

#### 3.3.1 Naive

Naive is an official baseline of CAFA. For given *P*_*j*_, the score that *P*_*j*_ is with *G*_*i*_ can be computed simply by the frequency of *G*_*i*_ in *D*, as follows (Note that this method gives the same score for all *P*_*j*_):

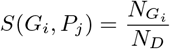

#### 3.3.2 BLAST-KNN

The idea follows a finding that using the similarity scores (bit-scores) between a query and its similar proteins improves the performance of using the sequence identity only [4]. For given *P*_*j*_, the score of BLAST-KNN *S*(*G*_*i*_, *P*_*j*_) is computed by first running BLAST to have the similarity score (bit-score) *B*(*P*_*j*_, *p*) between *P*_*j*_ and protein *p* to identify a set *H*_*j*_ of similar proteins to *P*_*j*_ in *D* using a certain cut-off value (set at *e*-value of 0.001 in our experiments) against *B*(*P*_j_, *p*) for all *p*. Finally the score can be obtained as follows:

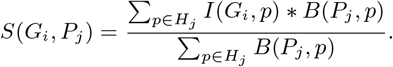

### 3.4 NetGO component: Net-KNN

Fig 2 shows the schematic procedure of Net-KNN for enumerating candidate GO terms for each protein by using network information. Given a test protein *P*_*j*_ and a protein network (from experiments), Net-KNN computes the score between *P*_*j*_ and GO term *G*_*i*_, *S*(*G*_*i*_, *P*_*j*_), through one of the following three ways and equation (1):

1) STRI: If *P*_*j*_ appears in STRING (meaning that *PV*_*j*_ exists), Net-KNN uses neighboring nodes *PV*_*k*_ of *PV*_*j*_ in the network;

2) HOMO: Net-KNN searches the most homologous protein in STRING to *P*_*j*_ by BLAST with cutoff E-value of 0.001 and uses this protein as *PV*_*j*_ and its neighboring nodes *PV*_*k*_;

3) NOHO: Net-KNN returns zero if no homologous proteins are found.

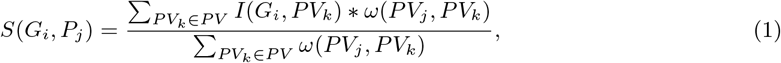

where *ω*(*PV*_*j*_, *PV*_*K*_) can be computed, assuming that *m* networks, *PN*^(*l*)^ (*l* =1,…, *m*), over the same set of nodes are given, as follows:

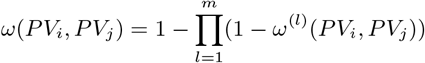

where again *ω*^(*l*)^ (*PV*_*i*_,*PV*_*j*_) is the confidence of the association between *PV*_*i*_, and *PV*_*j*_ in *PN*^(*l*)^ and *ω*(*PV*_*j*_, *PV*_*K*_) = 0 if there is no direct edges between *PV*_*i*_ and *PV*_*j*_.

**Figure 1:**
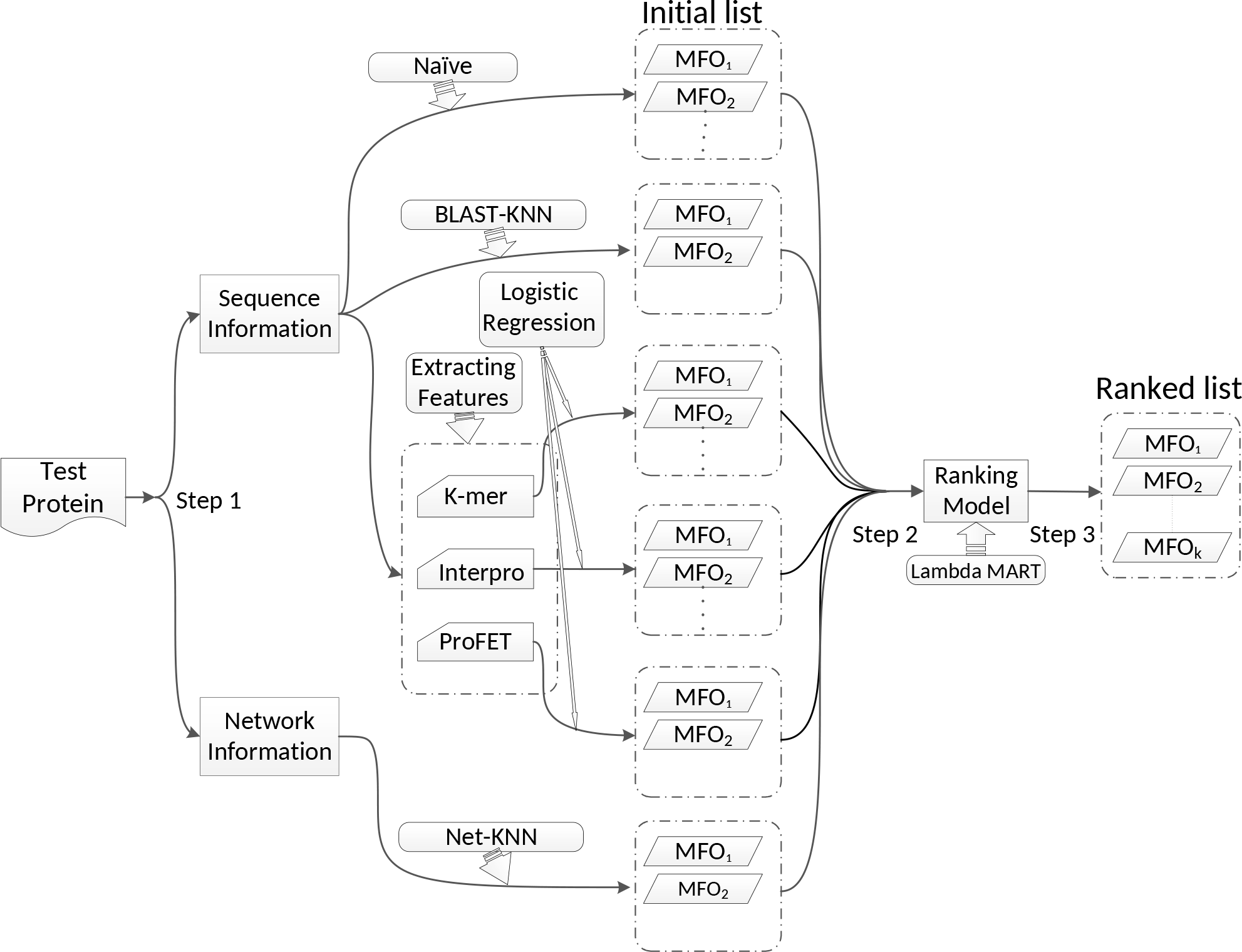
Entire scheme of NetGO with three steps for AFP. Sequence and network information are from five components and Net-KNN, respectively.

**Figure 2:**
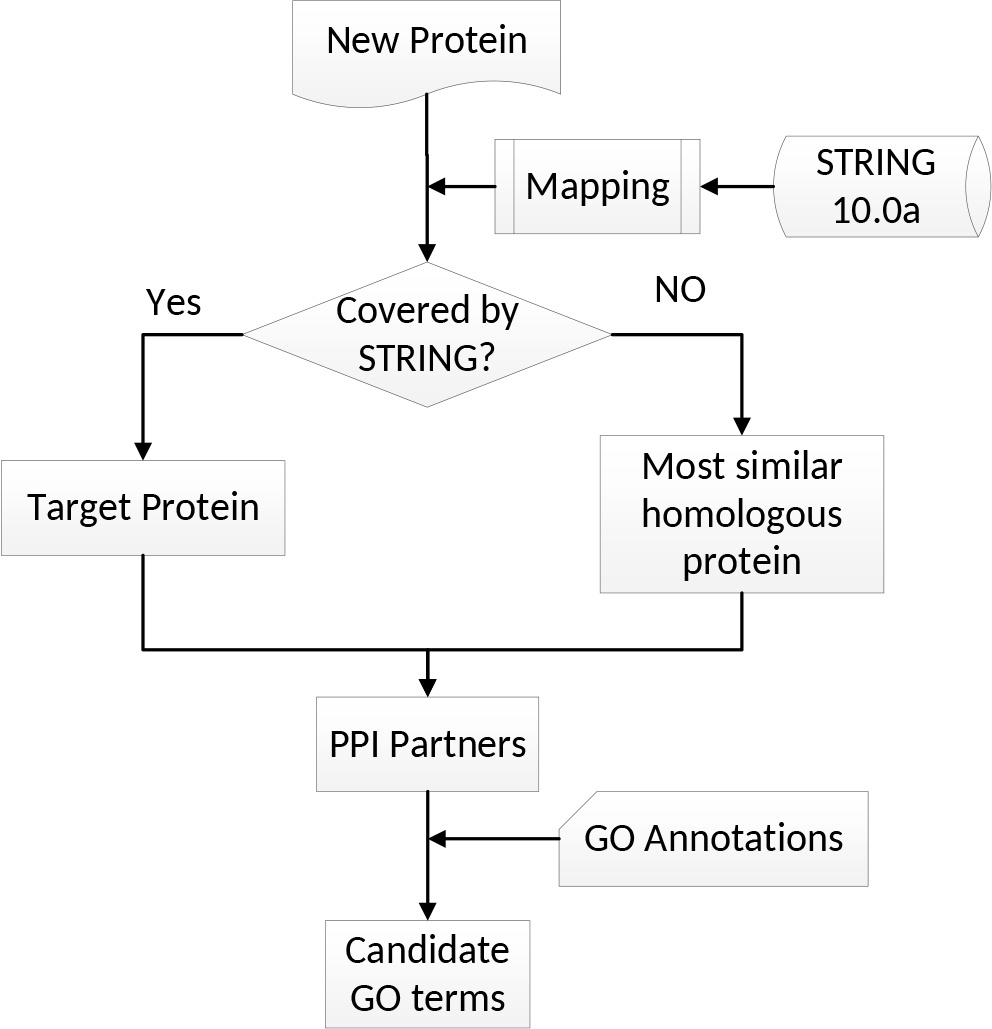
Procedure of Net-KNN.

The basic idea of Net-KNN is similar to that of BLAST-KNN, where the sequence similarity (bitscore by BLAST) in BLAST-KNN, and in Net-KNN is replaced by the association score (edge weight) in the network. Note that using different types of networks and their combinations changes the results and also final performance.

### 3.5 NetGO: Entire procedure

With three steps in total, NetGO starts by using six components that we have already explained.

#### Step 1: Generate candidates GO terms

Given a query protein, we run six component methods to produce the top-*k* GO terms from each component and then merge them to form the candidate GO terms (we used *k*=30 in our experiments. See Section 4.3). Note that reducing *k* is to focus on the most relevant GO terms to the query protein and also reduce the computational burden of the model.

#### Step 2: Generate features for ranking GO terms

We then generate features of the query protein by using the scores (of each of the candidate GO terms) predicted by all six component methods. As such, a six-dimensional feature vector is formed for each pair of a GO term and one query protein. All score values are between 0 and 1.

**Table 1:**
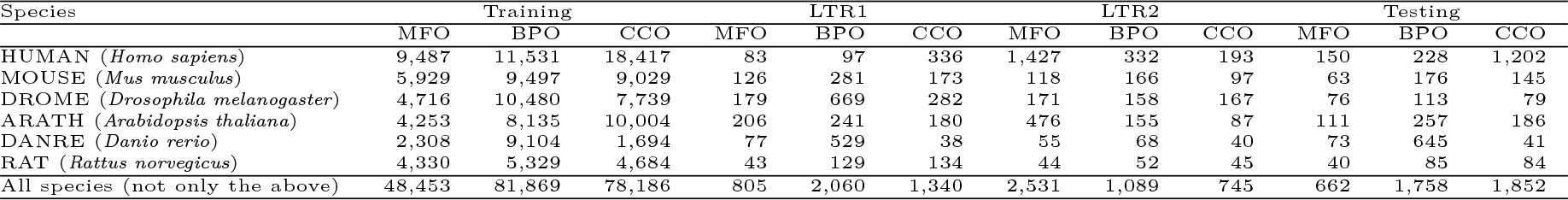
Data statistics (#proteins) on species with at least ten proteins for one domain of GO for each of the four datasets.

| Species | Training |  |  | LTR1 |  |  | LTR2 |  |  | Testing |  |  |
| --- | --- | --- | --- | --- | --- | --- | --- | --- | --- | --- | --- | --- |
|  | MFO | BPO | CCO | MFO | BPO | CCO | MFO | BPO | CCO | MFO | BPO | CCO |
| HUMAN (*Homo sapiens*) | 9,487 | 11,531 | 18,417 | 83 | 97 | 336 | 1,427 | 332 | 193 | 150 | 228 | 1,202 |
| MOUSE (*Mus musculus*) | 5,929 | 9,497 | 9,029 | 126 | 281 | 173 | 118 | 166 | 97 | 63 | 176 | 145 |
| DROME (*Drosophila melanogaster*) | 4,716 | 10,480 | 7,739 | 179 | 669 | 282 | 171 | 158 | 167 | 76 | 113 | 79 |
| ARATH (*Arabidopsis thaliana*) | 4,253 | 8,135 | 10,004 | 206 | 241 | 180 | 476 | 155 | 87 | 111 | 257 | 186 |
| DANRE (*Danio rerio*) | 2,308 | 9,104 | 1,694 | 77 | 529 | 38 | 55 | 68 | 40 | 73 | 645 | 41 |
| RAT (*Rattus norvegicus*) | 4,330 | 5,329 | 4,684 | 43 | 129 | 134 | 44 | 52 | 45 | 40 | 85 | 84 |
| All species (not only the above) | 48,453 | 81,869 | 78,186 | 805 | 2,060 | 1,340 | 2,531 | 1,089 | 745 | 662 | 1,758 | 1,852 |

**Table 2:** AUPR of Net-KNN using seven different types of networks: 0:neighbourhood to 5:database. “All” means the combination of all networks, while “All (binary, cut-off=0.5)” means the combination (union) of all networks, where each network edge weight (similarity) is transformed into a binary by using the cut-off value of 0.5

|  | All species |  |  | Eukaryote |  |  | Prokaryote |  |  |
| --- | --- | --- | --- | --- | --- | --- | --- | --- | --- |
|  | MFO | BPO | CCO | MFO | BPO | CCO | MFO | BPO | CCO |
| 0: neighbourhood | 0.047 | 0.023 | 0.195 | 0.052 | 0.230 | 0.195 | 0.030 | 0.025* | **0.197\*** |
| 1: fusion | 0.067 | 0.013 | 0.139 | 0.083 | 0.012 | 0.138 | 0.020 | 0.016 | 0.020 |
| 2: cooccurrence | 0.062 | 0.013 | 0.128 | 0.072 | 0.013 | 0.130 | 0.026 | 0.022 | 0.160 |
| 3: coexpression | 0.111 | 0.073* | 0.475 | 0.132 | 0.076* | 0.478 | 0.027 | 0.022 | 0.124 |
| 4: experiment | 0.101 | 0.066 | 0.504* | 0.125 | 0.071 | 0.509* | 0.033* | 0.020 | 0.023 |
| 5: database | 0.128* | 0.048 | 0.366 | 0.150* | 0.048 | 0.363 | 0.021 | 0.019 | 0.004 |
| All (binary, cut-off=0.5) | 0.119 | 0.053 | 0.481 | 0.143 | 0.056 | 0.482 | 0.028 | 0.015 | 0.134 |
| All (binary, cut-off=0.9) | 0.148 | 0.040 | 0.413 | 0.170 | 0.041 | 0.416 | **0.044** | 0.025 | 0.065 |
| All | **0.158** | **0.097** | **0.568** | **0.186** | **0.102** | **0.567** | 0.042 | **0.035** | 0.166 |

#### Step 3: Rank GO terms by learning to rank (LTR)

Finally we use LTR to rank all candidate GO terms of each query protein. All proteins in the training data and their candidate GO terms are used for training the LTR model. In this way, LTR allows to integrate multiple sequence- and network-based evidences of *no-knowledge* proteins.

### 3.6 Baseline methods

#### 3.6.1 Consensus and Consensus-BN

“Consensus” computes the score as follows:

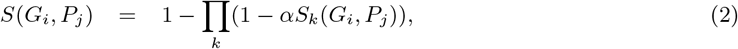

where *α* ∈ [0, 1] is an input constant to balance components by their importance where in our experiments, we used *α* = 1 that is most typical in many approaches such as COFACTOR and GAS-C for large-scale AFP[26, 28]. Also considering the idea of GAS-C, which used Consensus for BLAST and a network-based method only, we call Consensus of using BLAST-KNN and Net-KNN only “Consensus-BN”.

#### 3.6.2 DeepGO

We downloaded the code of DeepGO and trained the prediction model with the suggested parameters using the same training data as NetGO. Note that it only made predictions on the MFO, BPO and CCO terms that appeared more than 50, 250 and 50 times in the training data, respectively.

## 4 Experiments

### 4.1 Data

We collected data following CAFA1 [4], CAFA2 [5] and CAFA3:

1. Protein sequences We downloaded the FASTA-format files of all proteins from UniProt^4^ [3].
2. GO terms We first downloaded protein function annotation from SwissProt^5^ [7], GOA^6^ [31], and GO^7^ [2] in Oct. 2017, and then extracted all experimental annotations in: ‘EXP’, ‘IDA’, ‘IPI’, ‘IMP’, ‘IGI’, ‘IEP’, ‘TAS’, or ‘IC’, and then merged them to generate an annotation dataset (Note that SwissProt did not have annotation dates and so we downloaded data of SwissProt in Oct. 2015 and Oct. 2016).
3. Protein network We used version 10.0a of STRING[27], generated between Apr. 12, 2015 and Apr. 16, 2016, i.e. before T0 (Oct. 2016) of the experiment. This database covers 9,643,763 proteins from 2,031 organisms with 932,553,897 interactions in total. The networks of 359 organisms appearing in the training data were used in Net-KNN.

**Table 3:** Performance of NetGO, its component and competing methods.

|  | F<sub>max</sub> |  |  | AUPR |  |  |
| --- | --- | --- | --- | --- | --- | --- |
|  | MFO | BPO | CCO | MFO | BPO | CCO |
| Naive | 0.317 | 0.255 | 0.604 | 0.169 | 0.115 | 0.610 |
| BLAST-KNN | 0.589 | 0.283 | 0.641 | 0.453 | 0.110 | 0.560 |
| Net-KNN | 0.389 | 0.320 | 0.646 | 0.158 | 0.097 | 0.568 |
| DeepGO | 0.392 | 0.239 | 0.569 | 0.253 | 0.081 | 0.545 |
| Consensus-BN | 0.588 | 0.337 | 0.671 | 0.431 | 0.151 | 0.623 |
| Consensus | 0.576 | 0.327 | 0.670 | 0.443 | 0.146 | 0.651 |
| GOLabeler | 0.630 | 0.321 | 0.668 | 0.549 | 0.171 | 0.685 |
| NetGO(LTR1) | 0.629 | 0.338 | 0.669 | 0.546 | 0.191 | 0.697 |
| NetGO(LTR1+2) | **0.631** | **0.341** | **0.674** | **0.557** | **0.195** | **0.706** |

**Table 4:** In Testing, #proteins of STRI and HOMO in Net-KNN (see Sec.

| Species | MFO |  | BPO |  | CCO |  |
| --- | --- | --- | --- | --- | --- | --- |
|  | STRI | HOMO | STRI | HOMO | STRI | HOMO |
| HUMAN | 99 | 30 | 135 | 62 | 176 | 956 |
| MOUSE | 45 | 15 | 116 | 41 | 108 | 24 |
| DROME | 32 | 42 | 50 | 61 | 33 | 45 |
| ARATH | 76 | 25 | 168 | 64 | 125 | 46 |
| DANRE | 30 | 36 | 220 | 404 | 13 | 22 |
| RAT | 26 | 14 | 60 | 23 | 48 | 33 |
| All species | 399 | 184 | 889 | 690 | 559 | 1,148 |

**Table 5:** Performance by proteins in STRI and those in HOMO

|  | F<sub>max</sub> (by STRI) |  |  | AUPR (by STRI) |  |  | F<sub>max</sub> (by HOMO) |  |  | AUPR (by HOMO) |  |  |
| --- | --- | --- | --- | --- | --- | --- | --- | --- | --- | --- | --- | --- |
|  | MFO | BPO | CCO | MFO | BPO | CCO | MFO | BPO | CCO | MFO | BPO | CCO |
| Naive | 0.322 | 0.249 | 0.542 | 0.179 | 0.119 | 0.501 | 0.290 | 0.274 | 0.658 | 0.131 | 0.112 | 0.687 |
| BLAST-KNN | 0.608 | 0.310 | 0.597 | 0.470 | 0.137 | 0.486 | 0.627 | 0.267 | 0.690 | 0.529 | 0.099 | 0.619 |
| Net-KNN | 0.348 | 0.311 | 0.584 | 0.156 | 0.112 | 0.453 | 0.425 | **0.341** | 0.709 | 0.236 | 0.096 | 0.673 |
| GOLabeler | 0.631 | 0.333 | **0.626** | 0.554 | 0.188 | 0.617 | 0.665 | 0.312 | 0.715 | 0.596 | 0.152 | 0.736 |
| NetGO | **0.632** | **0.351** | 0.622 | **0.561** | **0.219** | **0.632** | **0.666** | 0.334 | **0.725** | **0.603** | **0.181** | **0.755** |

We then generated the four following datasets, which differ mainly in the time stamps when proteins are annotated.

1. Training: training for components All data annotated in Oct. 2015 or before.
2. LTR1: training for LTR *no-knowledge* proteins, experimentally annotated from Oct. 2015 to Oct. 2016 and not before Oct. 2015.
3. LTR2: training for LTR *limited-knowledge* proteins, experimentally annotated from Oct. 2015 to Oct. 2016 and not before Oct. 2015.
4. Testing: testing for competing methods All data experimentally annotated after Oct. 2016 by Oct. 2017 and not before Oct. 2016.

**Table 6:** Weighted node degree of proteins improved by NetGO

|  | MFO |  | BPO |  | CCO |  |
| --- | --- | --- | --- | --- | --- | --- |
|  | Degree | Num. | Degree | Num. | Degree | Num. |
| IMP | 13.72 | 264 | 27.12 | 915 | 51.51 | 950 |
| NOT | 15.32 | 204 | 22.73 | 649 | 34.63 | 531 |
| *p*-value | 0.502 |  | **0.013** |  | **8.326e-6** |  |

This time-series way of separating test data from training data is the same as CAFA. Also we used the same target species as CAFA3 in LTR1, LTR2 and Testing. Table 1 shows the number of proteins in the above four datasets. Note that the inputs of GOLabeler are sequences only, while the inputs of NetGO include both sequence and network information.

### 4.2 Performance Evaluation Measures

We used two measures for performance evaluation: AUPR (Area Under the Precision-Recall curve) and F_max_. AUPR is a standard evaluation metric in machine learning which punishes false positive prediction and so is suitable for highly imbalance data. F_max_ is an official metric of CAFA with the following definition.

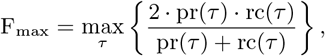

where pr(*τ*) and rc(*τ*) are *precision* and *recall*, respectively, obtained at some cut-off value, *τ*, defined as follows:

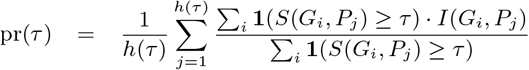

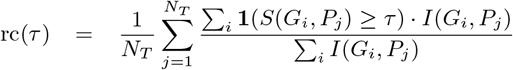

Given a test set of proteins, we first obtain the prediction score of each pair of a protein and a GO term. We then sort all pairs of proteins and GO terms according to the prediction scores, and evaluate the performance by F_max_ and AUPR. We used this AUPR for pairs all through experiments, except for the following two sections: In Section 4.4.4, AUPR per protein is computed by focusing on each protein, in which prediction scores of GO terms are computed, sorted and evaluated by AUPR. Similarly in Section 4.4.7, AUPR per GO term is computed for the prediction scores of proteins for each GO term.

### 4.3 Implementation and Parameter Settings

We processed the FASTA-format data by biopython^8^ and used sklearn^9^ to run logistic regression and xgboost [32] for LTR. Top 30 predictions from each component were merged, since this number provided the best performance in five-fold cross validation over LTR training data out of four values {10,30, 50 and 70} tested (see supplementary materials for the detailed results and other settings on components of GOLabeler and NetGO).

### 4.4 Results

To validate the performance of NetGO and other competing methods, we resampled the instances in testing with replacement 100 times (bootstrap with replacement) to make the experiment reliable. Except for the performance evaluation measures, we used the paired *t*-test to statistically evaluate the performance difference between the best performance (in boldface in tables) and all others. The result was considered significant if *p*-value was smaller than 0.05. In the following tables, the best performance value is underlined if the value is statistically significant (see the supplementary materials for detailed p-values).

**Table 7:** Species-specific performance: HUMAN and ARATH

|  | F<sub>max</sub> (HUMAN) |  |  | AUPR (HUMAN) |  |  | F<sub>max</sub> (ARATH) |  |  | AUPR (ARATH) |  |  |
| --- | --- | --- | --- | --- | --- | --- | --- | --- | --- | --- | --- | --- |
|  | MFO | BPO | CCO | MFO | BPO | CCO | MFO | BPO | CCO | MFO | BPO | CCO |
| Naive | 0.359 | 0.241 | 0.661 | 0.194 | 0.109 | 0.702 | 0.322 | 0.249 | 0.708 | 0.181 | 0.124 | 0.681 |
| BLAST-KNN | 0.560 | 0.349 | 0.670 | 0.356 | 0.142 | 0.593 | 0.625 | 0.318 | 0.648 | 0.531 | 0.132 | 0.529 |
| Net-KNN | 0.366 | 0.297 | 0.678 | 0.176 | 0.066 | 0.612 | 0.365 | 0.294 | 0.691 | 0.203 | 0.128 | 0.638 |
| GOLabeler | 0.618 | 0.375 | 0.703 | 0.502 | 0.216 | 0.726 | 0.702 | 0.345 | **0.735** | **0.701** | 0.213 | 0.784 |
| NetGO | **0.626** | **0.389** | **0.710** | **0.504** | **0.224** | **0.742** | **0.704** | **0.352** | **0.735** | 0.699 | **0.224** | **0.791** |

**Table 8:** Performance comparisons over difficult proteins

|  | F<sub>max</sub> |  |  | AUPR |  |  |
| --- | --- | --- | --- | --- | --- | --- |
|  | MFO | BPO | CCO | MFO | BPO | CCO |
| Naive | 0.317 | 0.247 | 0.545 | 0.175 | 0.116 | 0.513 |
| BLAST-KNN | 0.564 | 0.268 | 0.552 | 0.411 | 0.098 | 0.415 |
| Net-KNN | 0.334 | 0.284 | 0.566 | 0.155 | 0.082 | 0.405 |
| GOLabeler | **0.617** | 0.319 | 0.610 | 0.528 | 0.171 | 0.605 |
| NetGO | **0.617** | **0.336** | **0.613** | **0.537** | **0.190** | **0.623** |

#### 4.4. Performance of Net-KNN with different networks

Before validating the performance of NetGO, we examined AUPR of Net-KNN by using six different types of networks in STRING: 0:neighbourhood, 1:fusion, 2:co-occurrence, 3:coexpression, 4:experiment and 5:database. To avoid possible information leak, we didn’t use the sixth type of network (6:textmining) in STRING. Table 2 reports AUPR obtained by using each individual network (upper part) and by combining all of these networks (lower part). “All” in the table means the combination of all networks according to equation (1). In the upper part, the highest and second highest scores are marked with asterisk and displayed in italic, respectively.

We have three main findings: 1) Using all networks (“All” or “All(binary, cut-off=0.9)” in Table 2) achieved the best performance, except for CCO of prokaryote. 2) Among six networks, no one performed the best in more than three cases. It implies that the role of each network is different from each other. For example, 0:neighbourhood and 2:cooccurrence achieved the best and second for CCO of prokaryote, respectively. However, they are ranked 4th and 6th for CCO of eukaryote, respectively. This indicates that they are useful only for predicting prokaryote proteins. 3) Among three settings of network combination, keeping the association scores of network edge (“All” in Table 2) achieved the best performance in all nine cases, except MFO of Prokaryote. This implies that transforming the association scores into binary by using a cut-off value will cause information loss. So hereafter we use “All” as a component of NetGO.

#### 4.4.2 Performance of NetGO, its components and competing methods

Table 3 reports the performance results of NetGO, GOLabeler and other compared methods over Testing. In the upper part, we show the result of component methods: Naive, BLAST-KNN and Net-KNN. BLAST-KNN was the best for MFO, while Net-KNN was the best for BPO and CCO. In the middle part, we show the result of four competing methods, DeepGO, Consensus, Consensus-BN, and GOLabeler. The middle part indicates that NetGO outperformed four competing methods. In particular, NetGO performed better than GOLabeler and DeepGO significantly and constantly under all six cases. This implies that incorporating network information is useful for predictive performance improvement. Moreover, the under-performed DeepGO underlines the weakness of deep learning based methods that work on small number of GO terms due to insufficient training data and high computational complexity. Finally, in the lower part, we compared two variants of NetGO. NetGO(LTR1) used LTR1 only to train the ranking model, while NetGO(LTR1+2) used both LTR1 and LTR2. We can see that using both LTR1 and LTR2 achieved the best performance, especially in AUPR, which is consistent with the case in GOLabeler[11]. Hereafter, to make full use of massive network information, we show the results of NetGO by using both LTR1 and LTR2 that incorporates all 6 types of networks in STRING.

#### 4.4.3 Performance by proteins in STRI and in HOMO

In Net-KNN, in which we compute the association score between a protein and a GO term, proteins are first classified as STRI, HOMO, and NOHO (see Sec. 3.4). That is, STRI are proteins appearing in STRING, while HOMO are proteins being absent from STRING but having their homologous proteins appearing in STRING. Table 4 lists the number of proteins in STRI and HOMO in Testing, where the total number of proteins in Testing is given in Table 1. For example, Table 1 shows 1,758 proteins in BPO. Among them, 1,579 (89.8%) out of 1758 have network information, with 889 (50.6%) and 690 (39.2%) being in STRI and HOMO, respectively. For validating the performance of proteins in Testing, we divided them into those in STRI and those in HOMO. Table 5 reports the performance results. From the table, we can see that again NetGO achieved the best performance among the five competing methods under all of the settings, except for only two cases.

**Table 9:** AUPR of NetGO and other methods for four groups in Testing which are divided by #annotations per GO terms

|  | MFO |  |  |  | BPO |  |  |  | CCO |  |  |  |
| --- | --- | --- | --- | --- | --- | --- | --- | --- | --- | --- | --- | --- |
|  | >300 | 101-300 | 31-100 | 11-30 | >300 | 101-300 | 31-100 | 11-30 | >300 | 101-300 | 31-100 | 11-30 |
| BLAST-KNN | 0.839 | 0.643 | 0.555 | 0.538 | 0.460 | 0.210 | 0.129 | 0.105 | 0.723 | 0.325 | 0.186 | 0.139 |
| Net-KNN | 0.778 | 0.399 | 0.342 | 0.223 | **0.553** | 0.197 | 0.111 | 0.095 | 0.734 | 0.270 | 0.172 | 0.129 |
| GOLabeler | **0.911** | **0.717** | 0.595 | **0.585** | 0.466 | 0.222 | 0.151 | 0.110 | 0.732 | 0.362 | 0.209 | 0.137 |
| NetGO | 0.906 | 0.708 | **0.598** | 0.581 | 0.532 | **0.245** | **0.167** | **0.125** | **0.758** | **0.370** | **0.219** | **0.160** |

**Figure 3:**
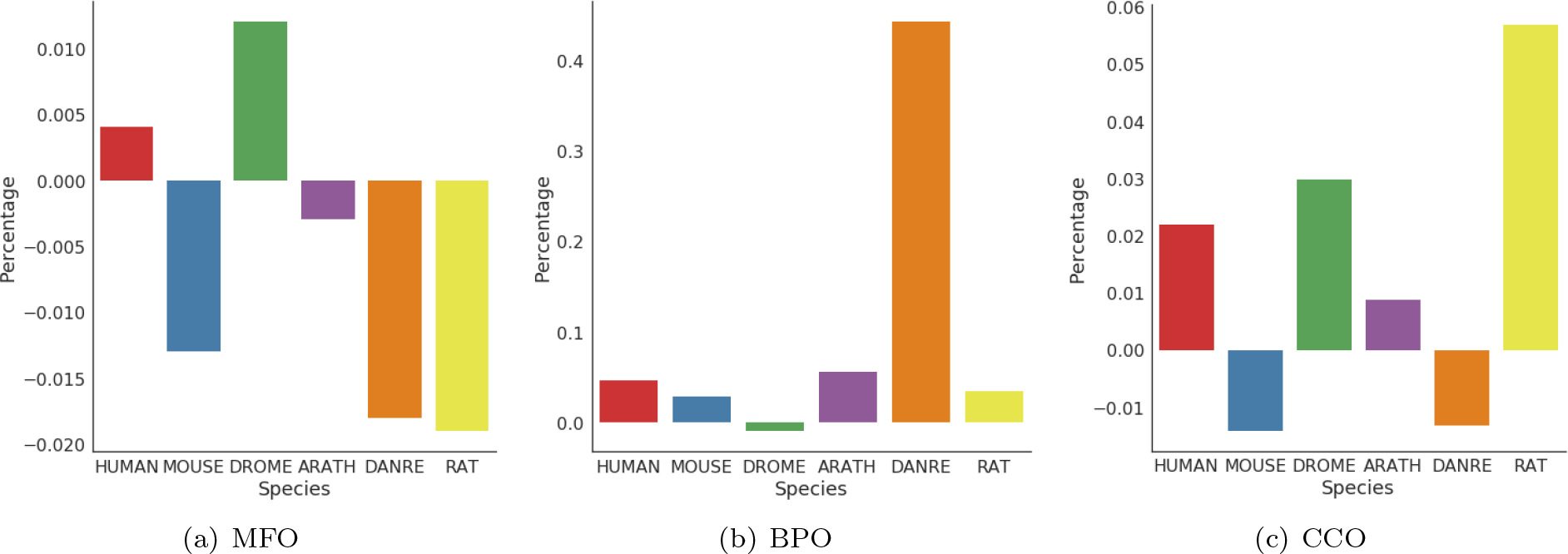
Improvement in AUPR of NetGO over GOLabeler

**Table 10:** Percentage of GO terms with highest AUC in each group

|  | MFO |  |  |  | BPO |  |  |  | CCO |  |  |  |
| --- | --- | --- | --- | --- | --- | --- | --- | --- | --- | --- | --- | --- |
|  | >300 | 101-300 | 31-100 | 11-30 | >300 | 101-300 | 31-100 | 11-30 | >300 | 101-300 | 31-100 | 11-30 |
| Total | 1 | 5 | 31 | 60 | 11 | 65 | 188 | 398 | 21 | 12 | 35 | 68 |
| GOLabeler | 1(100.0%) | 5(100.0%) | 17(54.8%) | 31(51.7%) | 1(9.1%) | 14(21.5%) | 52(27.7%) | 131(32.9%) | 10(47.6%) | 3(25.0%) | 15(42.9%) | 30(44.1%) |
| NetGO | 0(0.0%) | 0(0.0%) | 14(45.2%) | 24(40.0%) | 10(90.9%) | 51(78.5%) | 136(72.3%) | 264(66.3%) | 18(85.7%) | 9(75.0%) | 20(57.1%) | 38(55.9%) |

#### 4.4.4 Weighted node degree of performance enhanced proteins

We first computed AUPR per protein and then grouped all proteins in Testing into two subsets: 1) IMP:the performance was improved by NetGO and 2) NOT: not improved, according to AUPR. We then computed the weighted node degrees of those proteins in the network of STRING in this way: *PV*_*i*_ is the sum of *ω*(*PV*_*i*_, *PV*_*j*_) over all its neighbouring nodes *PV*_*j*_. If a given protein belongs to HOMO (see Sec. 3.4), we use the homologous protein with the highest bitscore by BLAST for *PV*_*i*_ instead. Also we evaluated the difference in weighted node degrees of two subsets by using *p*-values of *t*-test, assuming the normal distribution. Table 6 shows the average weighted node degree with *p*-values. It indicates that the IMP is always higher than NOT, with being statistically significant for BPO and CCO. This result implies that a higher node degree might be a key to improve the performance in BPO and CCO by using network information.

#### 4.4.5 Species-specific performance

In this experiment, we examined the individual performance of different species listed in Table 1. Table 7 reports the performance of HUMAN and ARATH, which have the top and second largest number of proteins, respectively. The table again shows that NetGO achieved the highest scores under all the settings, except for AUPR for MFO of ARATH. Also, we validated the performance improvement in AUPR of NetGO from GOLabeler on all six species. Figure 3 shows the improvement of AUPR for three domains of six species. The performance improvement was obvious for five out of six species in BPO and four out of six species in CCO, while that depends upon species for MFO. This result implies NetGO or network information is useful for BPO and CCO.

**Table 11:** Predicted GO terms of Q9BYW3 in BPO by NetGO and competing methods. Correctly predicted GO terms are in pink. B-K and N-K mean BLAST-KNN and Net-KNN, respectively. The last column shows the number of correctly predicted GO terms.

| Method | Predicted GO terms | C |
| --- | --- | --- |
| Naive | GO:0008150, GO:0009987, GO:0065007, GO:0050896, GO:0008152, GO:0032501, GO:0032502, GO:0071840, GO:0050789, GO:0071704, GO:0044237, GO:0048856, GO:0044238, GO:0007275, GO:0006807, GO:0016043, GO:0050794, GO:0048731 | 10 |
| B-K | GO:0008150, GO:0065007, GO:0032501, GO:0032502, GO:0009987, GO:0065008, GO:0009791, GO:0050789, GO:0009653, GO:0007568, GO:0019725, GO:0008340, GO:0007389, GO:0048856, GO:0007275, GO:0048646, GO:0042592, GO:0003002, GO:0009886, GO:0010259, GO:0045454, GO:0048513, GO:0048731, GO:0048645, GO:0050794, GO:0002164, GO:0009887, GO:0009790, GO:0048859, GO:0002165, GO:0035272, GO:0010160, GO:0002168, GO:0007436, GO:0022612, GO:0048732, GO:0007435, GO:0007431, GO:0007432 | 16 |
| N-K | GO:0008150, GO:0032502, GO:0032501, GO:0009987, GO:0065007, GO:0048856, GO:0007275, GO:0048731, GO:0048513 | 9 |
| DeepGO | GO:0008150, GO:0065007, GO:0050896, GO:0009987, GO:0022414, GO:0000003, GO:0050789, GO:0006950, GO:0006807, GO:0051716, GO:0044237, GO:0071704, GO:0044238, GO:0050794, GO:0034641, GO:0006139, GO:0046483, GO:0006725, GO:1901360, GO:0044260, GO:0019222, GO:0043170, GO:0048518, GO:0033554, GO:0048519, GO:0090304, GO:0006259, GO:0060255, GO:0080090, GO:0051171, GO:0031323, GO:0006974, GO:0048522, GO:0010468, GO:0031326, GO:0019219 | 4 |
| GOLabeler | GO:0008150, GO:0009987, GO:0065007, GO:0032501, GO:0032502, GO:0050896, GO:0008152, GO:0050789, GO:0048856, GO:0007275, GO:0065008, GO:0019725, GO:0007568, GO:0008340, GO:0042221, GO:0009653, GO:0050794, GO:0048731, GO:0042592, GO:0048513, GO:0002165, GO:0009886, GO:0002164, GO:0045454, GO:0010259, GO:0048859, GO:0048645, GO:0009790, GO:0009887, GO:0022612, GO:0002168, GO:0010160, GO:0035272, GO:0007436, GO:0048732, GO:0007432, GO:0007435, GO:0007431 | 16 |
| NetGO | GO:0008150, GO:0009987, GO:0065007, GO:0032501, GO:0032502, GO:0050896, GO:0008152, GO:0050789, GO:0048856, GO:0007275, GO:0065008, GO:0009653, GO:0050794, GO:0048731, GO:0048513, GO:0042592, GO:0009790, GO:0009887, GO:0048732 | 14 |
| Ground Truth | GO:0008150, GO:0065007, GO:0032501, GO:0009987, GO:0032502, GO:0019725, GO:0048856, GO:0007275, GO:0050789, GO:0065008, GO:0048513, GO:0050794, GO:0042592, GO:0035295, GO:0045454, GO:0048731, GO:0055123, GO:0048732, GO:0048565, GO:0031016, GO:0061008, GO:0001889 |  |

#### 4.4.6 Prediction on difficult proteins

Proteins with BLAST identity of less than 0.6 to any protein in training data are called “difficult proteins” [5]. It is hard to infer function of difficult proteins through homology, which makes annotating difficult proteins a challenging problem. Table 8 shows the performance obtained by applying the five compared methods to the difficult proteins. NetGO achieved the best performance under all six settings, implying that NetGO is most reliable and robust method among the compared methods.

#### 4.4.7 Comparison over groups divided by #annotations per GO term

According to the number of annotations per GO term: >300, 101-300, 31-100 and 11-30, we grouped annotations (GO terms) in Testing into four groups. Table 9 lists AUPR computed in each group. Obviously, NetGO outperformed other methods in all 12 settings except for four cases. The number (and percentage) of GO terms with the highest AUC in each setting is given in Table 10. It is clear that NetGO outperformed GOLabeler for BPO and CCO, while the two methods are comparable for MFO.

#### 4.4.8 Case study:*β*-defensin 126 (Q9BYW3)

By a typical real example, we finally illustrate the real performance difference on annotating GO to unknown proteins by using NetGO and other methods. Table 11 lists the predicted GO terms of BPO for Q9BYW3, with 18 true GO terms (the bottom row). Fig 4 plots the directed acyclic graph with these 18 GO terms. Fig 4 shows the directed acyclic graph with these 18 GO terms. As a difficult protein, Q9BYW3 has no homologous proteins (cut-off e-value at 0.001). For Q9BYW3, BLAST-KNN could not predict any GO terms. Naive predicted 12 GO terms, with only two true GO terms. Net-KNN detected 12 true GO terms out of its predicted 23 GO terms. DeepGO predicted 3 out of 14, while GOLabeler predicted 4 out of 9. Finally, NetGO correctly predicted 12 out of 17. The highest number of GO terms by both Net-KNN and NetGO is 12, with the error being only five by NetGO but eleven by Net-KNN. From this real example, this result demonstrates the high predictive performance of NetGO over other competing methods.

**Figure 4:**
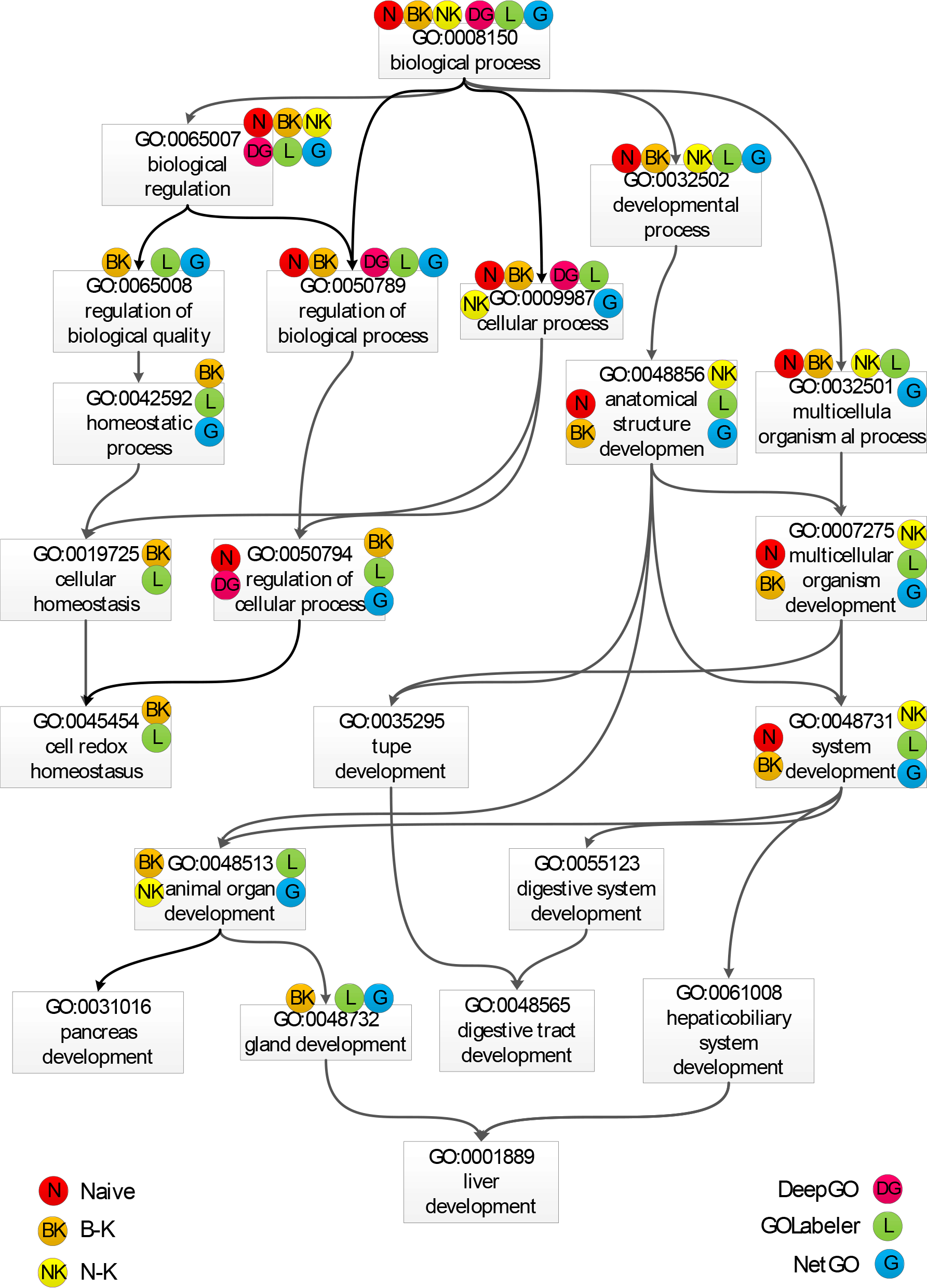
Predicted GO terms of Q9BYW3 in DAG of BPO by different methods.

## 5 Conclusion and discussion

The large-scale AFP has become an imperative issue, due to the explosive growth of protein sequence. The-state-of-the-art method, GOLabeler, uses only sequences information. Specifically, GOLabeler integrates multiple types of evidence, such as homology, domain, family, and motif in the powerful LTR framework. In addition, several approaches, such as MS-kNN, COFACTOR, GAS-C and DeepGO, have attempted to incorporate network-based information for AFP in the CAFA setting with limited success.

In this paper, we have present NetGO that incorporates massive network information. Extensive experiments show that NetGO outperformed GOLabeler and DeepGO significantly in all three domains, especially BPO and CCO, under the CAFA data setting. The reasons for such performance of NetGO are threefold: 1) the powerful LTR integration framework; 2) massive and complete network information from STRING; and 3) various sequence information. Many previous work report that interacting proteins tend to share the same GO function in BPO and CCO [26, 17, 16]. This is confirmed by our studies, in which the performance of NetGO significantly improved GOLabeler under these two domains, especially in BPO. Also a high-degree node (protein) tends to be associated with more GO terms, which means that high-degree proteins are more important in AFP [19]. NetGO uses confidence of associations instead of binary relations to compute node degree. This finding may partially explain the success of NetGO. The number of proteins with improved prediction performance is bigger than that of those with reduced performance. Interestingly, the proteins with improved prediction performance have higher degrees than those with reduced performance in BPO and CCO. This implies that NetGO performs better for high degree proteins due to its better performance for these proteins. Furthermore, we grouped GO terms due to the number of annotations per GO term. As such, the performance is improved over all of the groups. This fact means that the improvement was not biased to generic (major) GO terms. This highlights the advantage of NetGO in predicting both generic and more specific GO terms.

We have seen the various performance improvement by NetGO over GOLabeler under different settings, such as species centric evaluation, and difficult proteins. However there would be still challenges in the future. One is to annotate proteins of minor organism with highly less annotations. Another interesting future work will develop a more advanced network-based component than Net-KNN.

## Footnotes

1 http://biofunctionprediction.org/cafa/

2 http://biofunctionprediction.org/meetings/2017-meeting/

3 http://www.ebi.ac.uk/interpro/interproscan.html

4 http://www.uniprot.org/downloads

5 http://www.uniprot.org/downloads

6 http://www.ebi.ac.uk/GOA

7 http://geneontology.org/page/download-annotations

8 http://biopython.org/

9 http://scikit-learn.org/stable/index.html

